# A model for angiogenesis suppression via SERPINF1 in the surrounding pro-acinar microenvironment during human fetal pancreas development

**DOI:** 10.1101/2023.10.25.559143

**Authors:** Pratik Nailesh Mehta, Charles Giardina

## Abstract

Pancreas hormone-producing endocrine cell islets are integrated with the blood supply, whereas pancreatic digestive-enzyme producing acini are not. The functional result is that pancreatic acinar cells release their digestive enzymes into the ductal system and the duodenum to aid in digestion, while pancreatic hormone-producing endocrine cells release their hormones, such as insulin and glucagon, into the bloodstream. Despite this understanding, the mechanism for how acinar cells are free of vascularization is not completely clear. Based on an analysis of publicly available single cell RNA-sequencing data (GSE110154) of the human fetal pancreas during gestational weeks 12 through 22, a model is proposed that draws upon the anti-angiogenic properties of SERPINF1. This model envisions +PTF1A/+SOX9/+CPA1 pancreatic pro-acinar tip progenitor and +COL1A2/+PDGFRA pancreatic stromal fibroblast cell populations that surround the developing acini secreting SERPINF1 (PEDF) to inhibit angiogenesis and endothelial cell generation. This secretion could maintain an acinar microenvironment that is free from robust vascularization and integration of blood vessels as seen in mature pancreatic islet of Langerhans/endocrine cells.

## 1.1 Introduction

Pancreatic diseases such as insulin-dependent diabetes and pancreatic cancer could be cured by discovering how to generate human pancreases in vitro using stem cells for transplantation. To this end, enhancing the current understanding of the various components of pancreas developmental biology will enable the field to improve differentiation protocols to work towards the goal of pancreas generation. A component that is well-known in the field is the integration of hormone-producing endocrine cell islets with the blood supply. Conversely, pancreatic digestive-enzyme producing acini are not integrated with the blood supply. The functional result of this relationship is that pancreatic acinar cells release their digestive enzymes into the ductal system and duodenum to aid in digestion, while pancreatic hormone-producing endocrine cells release their hormones, such as insulin and glucagon, into the blood to regulate blood glucose levels (as well as a range of other functions). Despite this understanding, the mechanism for how acinar cells are free from vascularization is not yet completely clear.

Using computational methods to analyze publicly available single cell RNA-sequencing data (GSE110154) of the human fetal pancreas during gestational weeks 12 through 22, Serine Protease Inhibitor Family 1 (SERPINF1), also known as Pigment Epithelium-Derived Factor (PEDF), was found to be expressed within a subset of pancreatic pro-acinar tip progenitors triple-positive for SRY-Box Transcription Factor 9 (SOX9), Pancreas Associated Transcription Factor 1a (PTF1A), and Carboxypeptidase A1 (CPA1) transcripts. A subpopulation of Collagen Type I Alpha 2 Chain (COL1A2) expressing pancreatic fibroblasts were also positive for SERPINF1/PEDF transcripts. Furthermore, within this double-positive +COL1A2/+SERPINF1 fibroblast population, there was a Platelet-Derived Growth Factor Receptor Alpha (PDGFRA) positive subset. SERPINF1/PEDF has been classified as a multifunctional protein with anti-angiogenic, anti-tumorigenic, and neurotropic properties^1,2^. Leveraging evidence from this analysis, a model is proposed that draws upon <u>the anti-angiogenic property of SERPINF1 which suggests that +PTF1A/+SOX9/+CPA1 pancreatic pro-acinar tip progenitor cells and +COL1A2/+PDGFRA pancreatic stromal fibroblasts surrounding the developing acini secrete SERPINF1 (PEDF) to inhibit angiogenesis/endothelial cell generation thereby maintaining an acinar microenvironment that is free from robust vascularization and integration of blood vessels in, contrast of what is seen pancreatic islet of Langerhans/endocrine cells.</u> (Figure 1.1)

**Figure 1.1:**
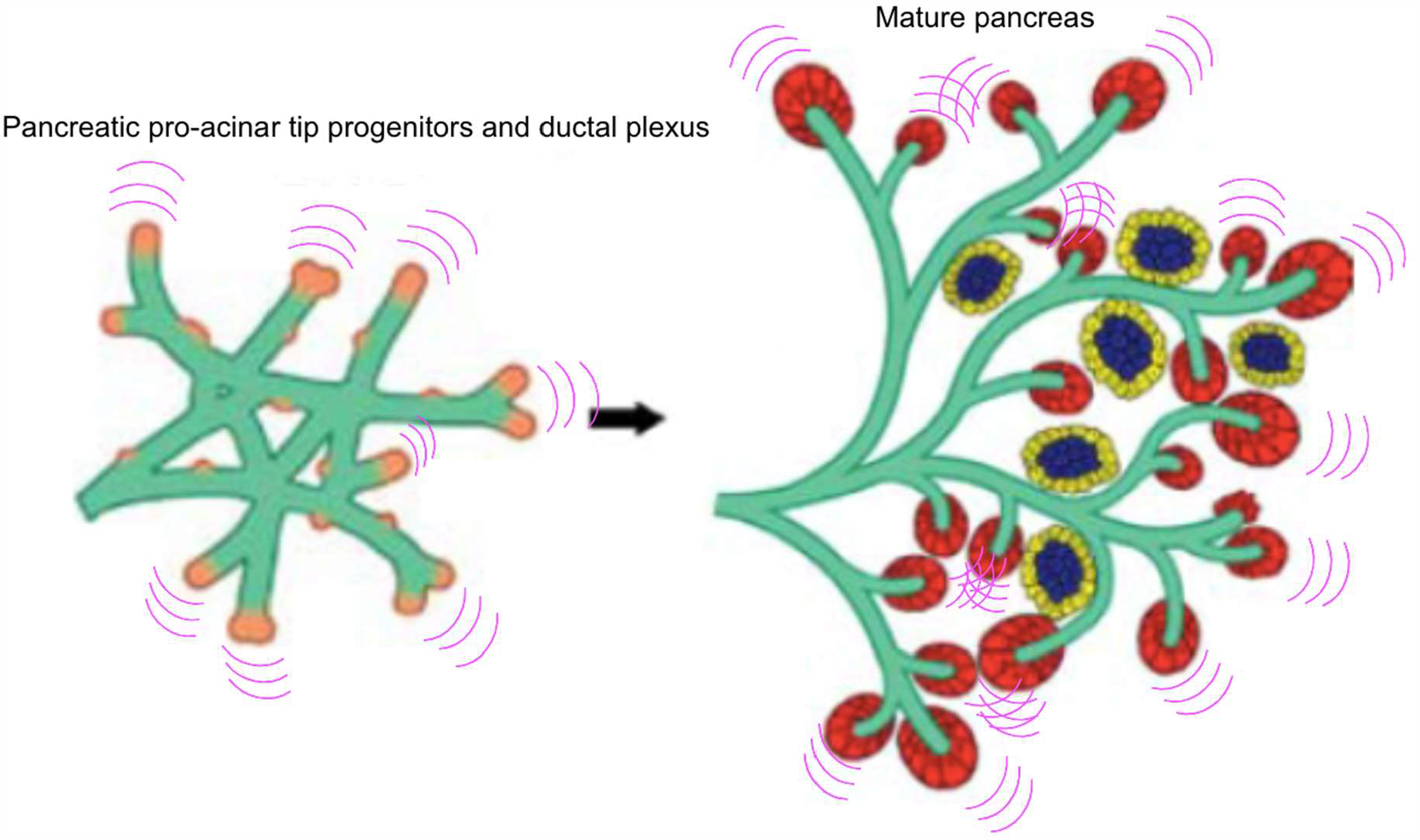
Image obtained and adapted from Pan and Wright^3^. Model for secretion of anti-angiogenic SERPINF1/PEDF in developing pancreas (left) and mature pancreas (right). Pink waves indicate secretion of SERPINF1/PEDF from pancreatic pro-acinar tip progenitors in orange (left) and from mature acini in red (right). Right: blue group of cells outlined in yellow cells are Islets of Langerhans or pancreatic endocrine cells

**Table 1.1:**
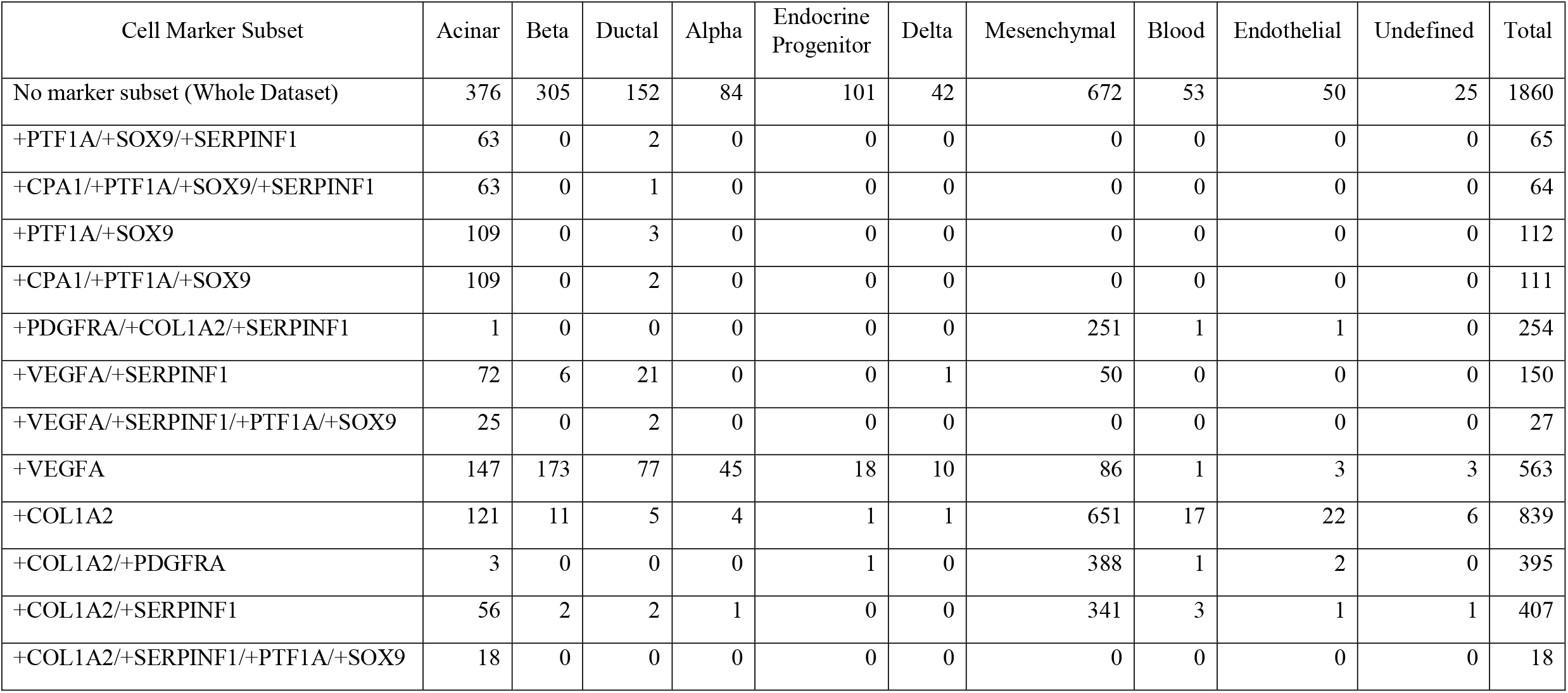
Cell counts grouped by cell-type using genes to subset data (+ indicates expression greater than 0 for each marker in Cell Marker Subset)

## 1.1 Methods

### 1.1.1 Data acquisition

The scRNA-seq data used for this analysis comes from the NCBI Gene Expression Omnibus (GEO: GSE110154), and I am thankful to Dr. Martin Enge and Dr. Efsun Arda for contributing this data publicly to the NCBI GEO. Dr. Martin Enge provided permission to use this data via email correspondence.

### 1.1.2 Data preparation and quality control

Individual cell raw read values were obtained from the uncompressed GSE110154_RAW.tar downloaded from the “Supplementary file” section on the GEO. Metadata was parsed out of the uncompressed series matrix file (GSE110154_series_matrix.txt.gz) obtained from the “Download family” section on the GEO. Bash and R scripting was used to combine the individual cell raw read values into a gene expression matrix and also generate a metadata dataframe to input into Seurat V4^4^. To reduce the number of genes that were most likely not involved in differentiating groups of cells, I did not change the default “min.cells = 3” parameter of CreateSeuratObject. In essence, this removed all genes that were expressed in less than 3 cells from the whole dataset. Likewise, the default “min.features = 200” parameter was not changed to serve as a way to remove dead cells or potentially empty wells. Setting this parameter to 200 removed all cells that had less than 200 genes detected (gene raw read values above 0). The Seurat object was generated: CreateSeuratObject(min.cells = 3, min.features = 200). The percentage of transcripts that map to mitochondrial genes for each cell was calculated and stored within the Seurat object: PercentageFeatureSet(pattern = “^MT”). The Seurat object was then normalized using the SCTransform^5^ function in conjunction with glmGamPoi^6^ within Seurat V4^4^ with parameters to regress out the percent of mitochondrial gene mapping: SCTransform(method = “glmGamPoi”, vars.to.regress = “percent.mt”). Principal components were generated to reduce the high-dimensionality of the scRNA-seq data using the variable features determined through SCTransform() for use in the successive steps of the workflow: RunPCA(features = VariableFeatures(), npcs = 100, approx=FALSE). The cells were then clustered using 100 principal components and the Leiden algorithm^7^.

### 1.1.3 Utilization of Seurat FeaturePlot function

The clustering resolution parameter was optimized based on the gene expression localization of the major pancreatic endocrine cell-type markers: FindNeighbors(graph.name = c(“NN”, “SNN”), dims = 1:100), FindClusters(graph.name = “SNN”, resolution = 2.9, algorithm = 4, group.singletons = TRUE). The default Seurat UMAP parameters were also modified to allow visually distinguishable cells and cell clusters: RunUMAP(min.dist = 1.3, n.neighbors = 90, dims = 1:100). The generic FeaturePlot command used to visualize gene expression was: FeaturePlot(reduction = “umap”, features = “my.favorite.gene”). To visualize gene-gene co-expression, two genes were added to the command and the “blend” option was enabled within the FeaturePlot function as such: FeaturePlot(reduction = “umap”, features = c(“my.first.favorite.gene”, “my.second.favorite.gene”), blend = TRUE).

### 1.1.4 General differential gene expression analysis for heatmap visualization

Differential gene expression (DGE) analysis was required to generate the data required for input in heatmap visualization. For DGE analysis, the Satija Lab (authors of Seurat) recommend using data normalized by following the standard Seurat normalization steps in the guided clustering workflow^8^. It is generally advised against using normalized data from SCTransform for DGE analysis. Data normalized through SCTransform is only superior for clustering cell types. After normalizing the data using the guided clustering workflow^8^ the following generic functions were used for differential gene expression analysis and heatmap generation: FindAllMarkers(assay = “RNA”, only.pos = TRUE) and DoHeatmap(assay = “RNA”).

### 1.1.5 Developmental signaling pathway—gene association screen

After a list of top differentially expressed genes per cluster was obtained using the FindAllMarkers function within Seurat, bash scripting was utilized to download gene-pathway associations from the Comparative Toxicogenomics Database. From the database file, the script extracted genes associated with developmental signaling pathways using signaling pathway keywords such as: “Wnt”, “TGF”, “Notch”, “Hedgehog”, “FGF”, so on and so forth. Using these genes, the script then searched the initial list of top differentially expressed genes per cluster for hits. The gene expression of these hits were mass-visualized using the Seurat FeaturePlot function. FeaturePlot hits that appeared to be expressed in a cell-type specific manner were stored, and one-by-one searched in the scientific literature to find novel connections to human pancreas development. One of the hits was SERPINF1 (PEDF).

## 1.2 Results

### 1.2.1 Endocrine cell markers used as “anchors” for optimal cell clustering

**Figure 1.2.1.1:**
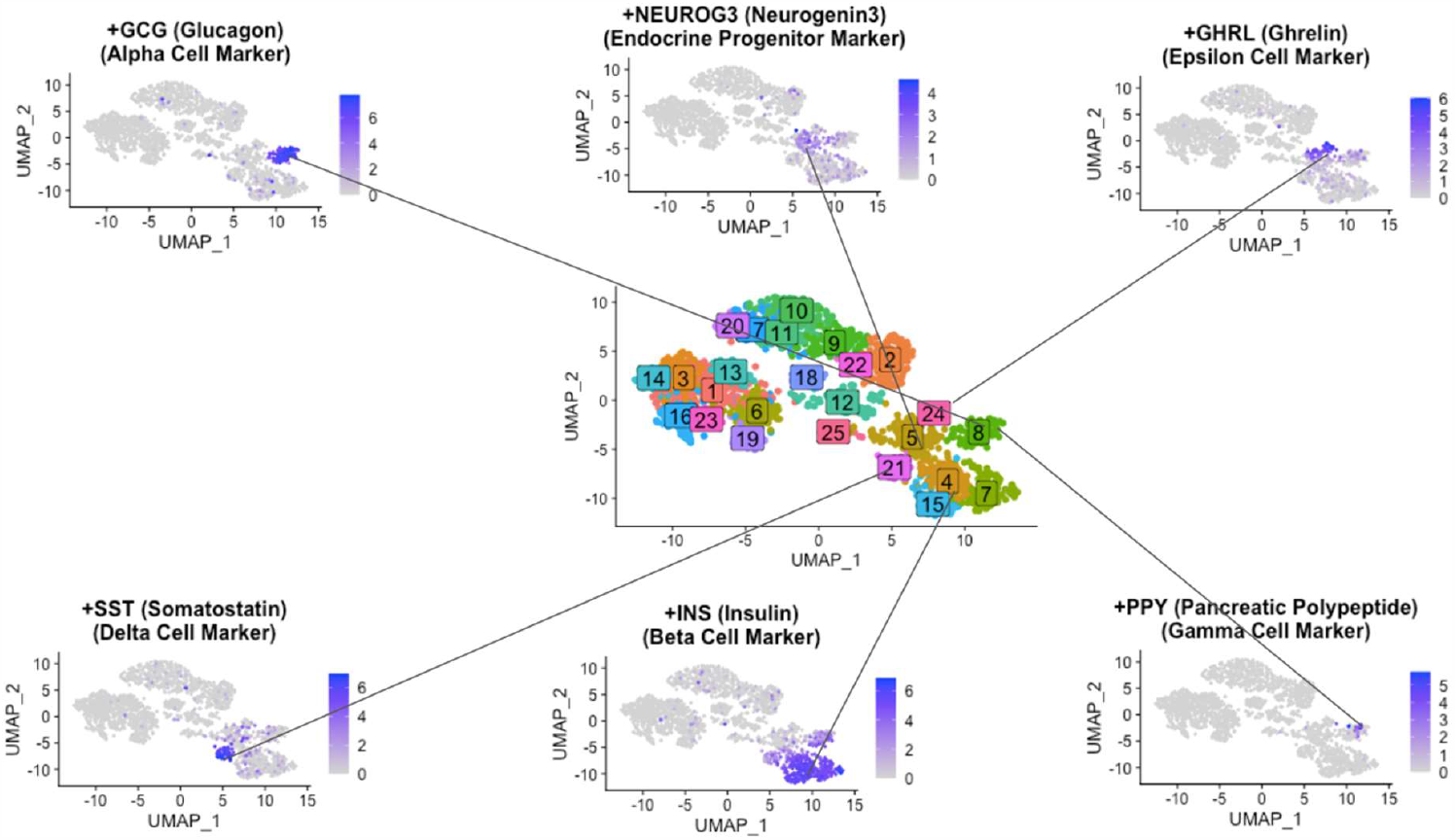
Middle: Multi-colored UMAP scatterplot of optimized cell clustering with colored boxed numbers indicating cluster number. Top three from left to right: Glucagon-expressing alpha cells residing within cluster 8, Neurogenin3-expression -marker for endocrine progenitors residing within cluster 5, Grehlin-expressing epsilon cells residing within cluster 24. Bottom three from left to right: Somatostatin-expressing delta cells residing within cluster 21, Insulin-expressing beta cells residing within clusters 4, 7 and 15, Pancreatic polypeptide-expressing cells also residing within cluster 8

To optimize cell-type clustering, the FeaturePlot function served as a key to visualize the gene expression of each endocrine cell-type marker and to compare it to the clustering that was generated for each adjustment of the resolution parameter. Simply, this was a “guess-and-check” method to see which resolution resulted in cell clustering that aligned with biologically-relevant pancreatic endocrine cell-type marker gene expression. Figure 1.2.1.1 below shows high-expression of somatostatin (SST) as a marker for endocrine delta cells localized to cluster 21 and high-expression of glucagon (GCG) as a marker for endocrine alpha cells localized to cluster 8. Insulin-expressing (INS) beta cells grouped in three subclusters (clusters 4, 7 and 15) with the optimized parameters that were selected. Neurogenin3 (NEUROG3), a marker of endocrine progenitors, was diffusely expressed in cluster 5. The primary endocrine cell-type marker and cluster alignment that was used to establish the clustering resolution for this analysis was ghrelin-expressing (GHRL) epsilon cells (cluster 24). Pancreatic polypeptide (PPY) expression, an endocrine gamma cell marker, was not suitable to use for cluster optimization because relative to the dataset the number of cells expressing the PPY gene was very low. Adjusting the clustering resolution or other parameters until the clusters became the size of the tiny population of PPY-expressing gamma cells would have resulted in, both, an unmanageable number of clusters and clusters that do not necessarily reveal anything biologically meaningful. Cluster 12, presumably endocrine progenitors, being positioned at the center of the clusters expressing differentiated endocrine cell-type markers was a sign of successful clustering. The central position of the endocrine progenitor population relative to the differentiated endocrine cell-types potentially provides an opportunity for analyzing the gene expression of cell lineage differentiation programs (single-cell trajectory analysis) [not discussed]. Interestingly, a +FEV^HI^ subpopulation of endocrine progenitor cells^9^ was identified in this dataset [not shown].

The optimal clustering resolution parameter was further validated using a heatmap to generally ensure that each cluster had a unique gene expression signature (Figure 1.2.1.2). In this case, cluster-specific unique gene expression signatures were verified by looking for unique high gene expression “blocks” (yellow) progressively “stepping down” diagonally from the top left of the heatmap to the bottom right. Either, each cluster had unique top differentially expressed genes that are upregulated at higher levels than other clusters or groups of clusters that shared the same top differentially expressed genes with unique levels of expression within those differentially expressed genes.

**Figure 1.2.1.2:**
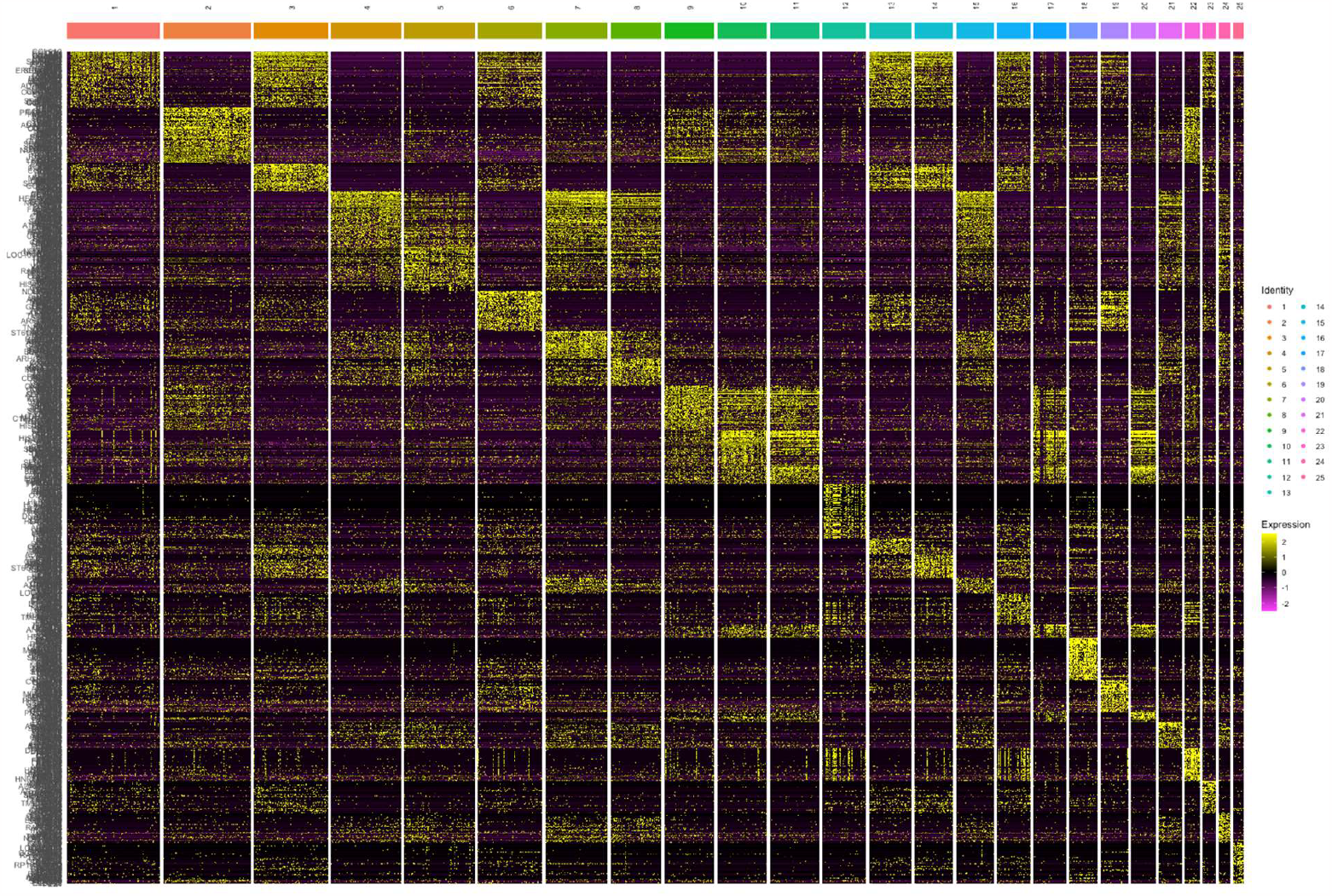
Heatmap of clusters. Genes on rows, cells grouped together into clusters on columns.

### 1.2.2 SERPINF1/PEDF expression in pro-acinar cells and stromal fibroblasts

To visualize gene-gene co-expression of SERPINF1 (PEDF) compared to other genes of interest in UMAP space, the FeaturePlot function of Seurat was used. FeaturePlots allowed the visualization of the relative gene expression level of individual cells across the entire dataset. Carboxypeptidase A1 (CPA1) and SERPINF1 (PEDF) displayed high-levels of co-expression in acinar cells (Figure 1.2.2.1). To verify that these +CPA1/+SERPINF1 (PEDF) cells also expressed other acinar cell markers, FeaturePlots were generated for other pancreatic acinar digestive enzymes (SPINK1, PRSS1, and CEL) (Figure 1.2.2.2). The Collagen Type I Alpha 2 Chain (COL1A2) fibroblast marker and SERPINF1/PEDF gene-gene co-expression FeaturePlot displayed a number of cells co-expressing both genes (Figure 1.2.2.3). Again to verify that these +COL1A2/+SERPINF1 (PEDF) cells also expressed other fibroblast cell markers, FeaturePlots were generated for other common pancreatic fibroblast genes (VIM, COL5A1, PDGFRA, PDGFRB) (Figure 1.2.2.4). PDGFRA being a common marker to subclassify fibroblasts, PDGFRA and SERPINF1/PEDF gene-gene co-expression FeaturePlot was generated to reveal PDGFRA as another positive marker for +SERPINF1/+PEDF fibroblasts (Figure 1.2.2.5). Because SERPINF1/PEDF has anti-angiogenic properties, the last gene-gene co-expression FeaturePlot pair of interest would include VEGFA with pro-angiogenic properties (Figure 1.2.2.6). Finally, to determine the number of cells expressing various markers combinations of interest, Table 1.1 was made.

**Figure 1.2.2.1:**
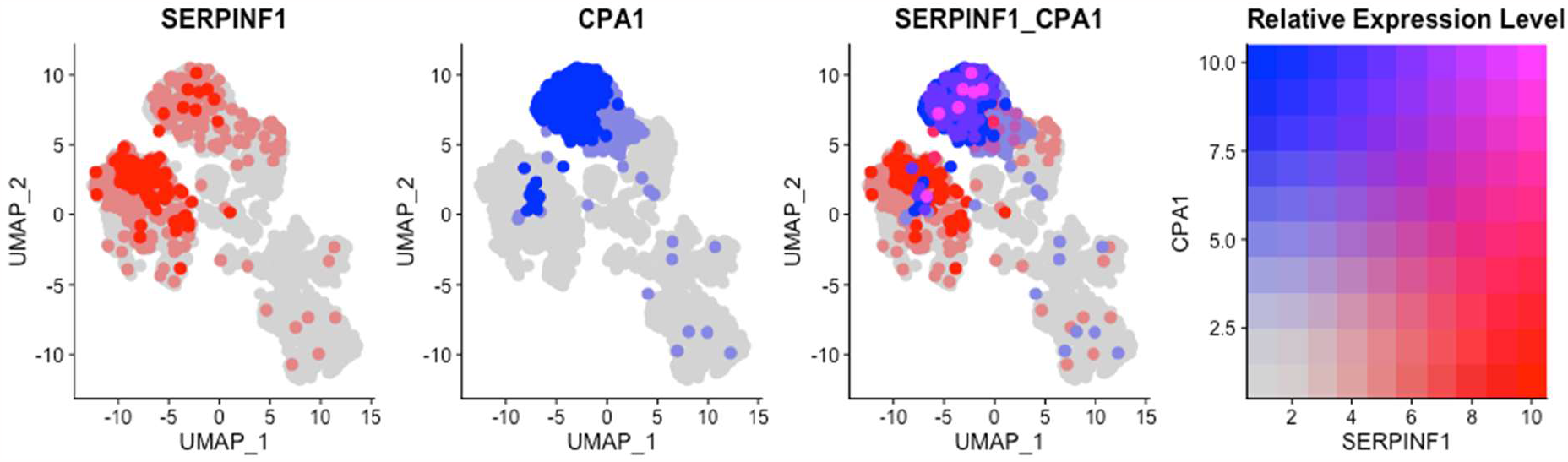
SERPINF1/PEDF and CPA1 (acinar cell marker) gene co-expression FeaturePlots. Far left to far right: Red SERPINF1/PEDF relative gene expression level, Blue CPA1 relative gene expression level, Red SERPINF1/PEDF and blue CPA1 merged (purple) relative gene expression level, Key of relative gene expression level for preceding three FeaturePlots.

**Figure 1.2.2.2:**
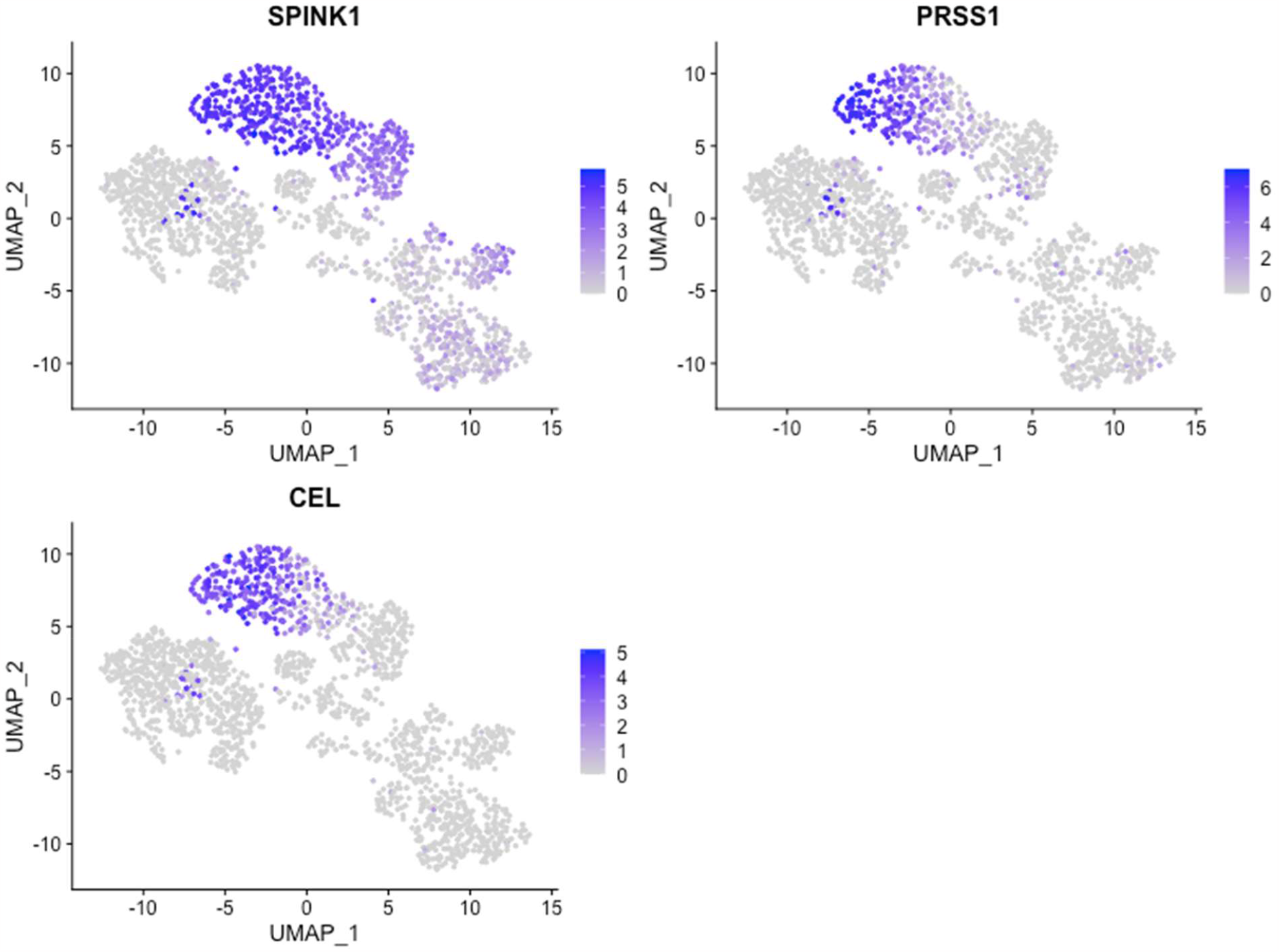
FeaturePlots of gene expression for pancreatic acinar digestive enzymes (SPINK1, PRSS1, and CEL)

**Figure 1.2.2.3:**
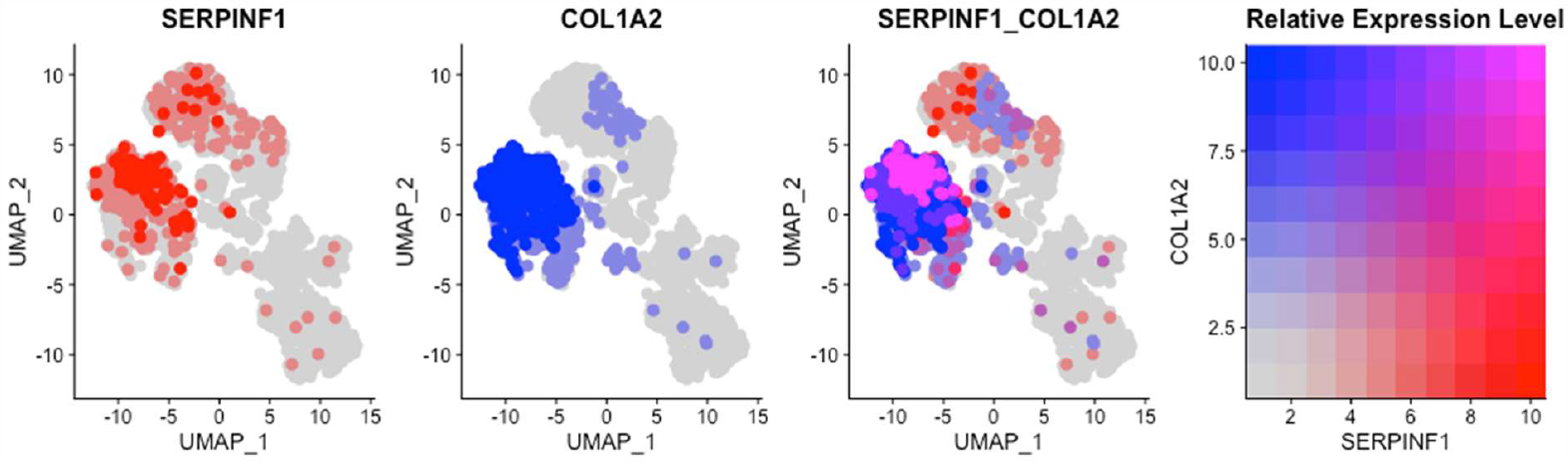
SERPINF1/PEDF and COL1A2 (fibroblast cell marker) gene co-expression FeaturePlots. Far left to far right: Red SERPINF1/PEDF relative gene expression level, Blue COL1A2 relative gene expression level, Red SERPINF1/PEDF and blue COL1A2 merged (purple) relative gene expression level, Key of relative gene expression level for preceding three FeaturePlots.

**Figure 1.2.2.4:**
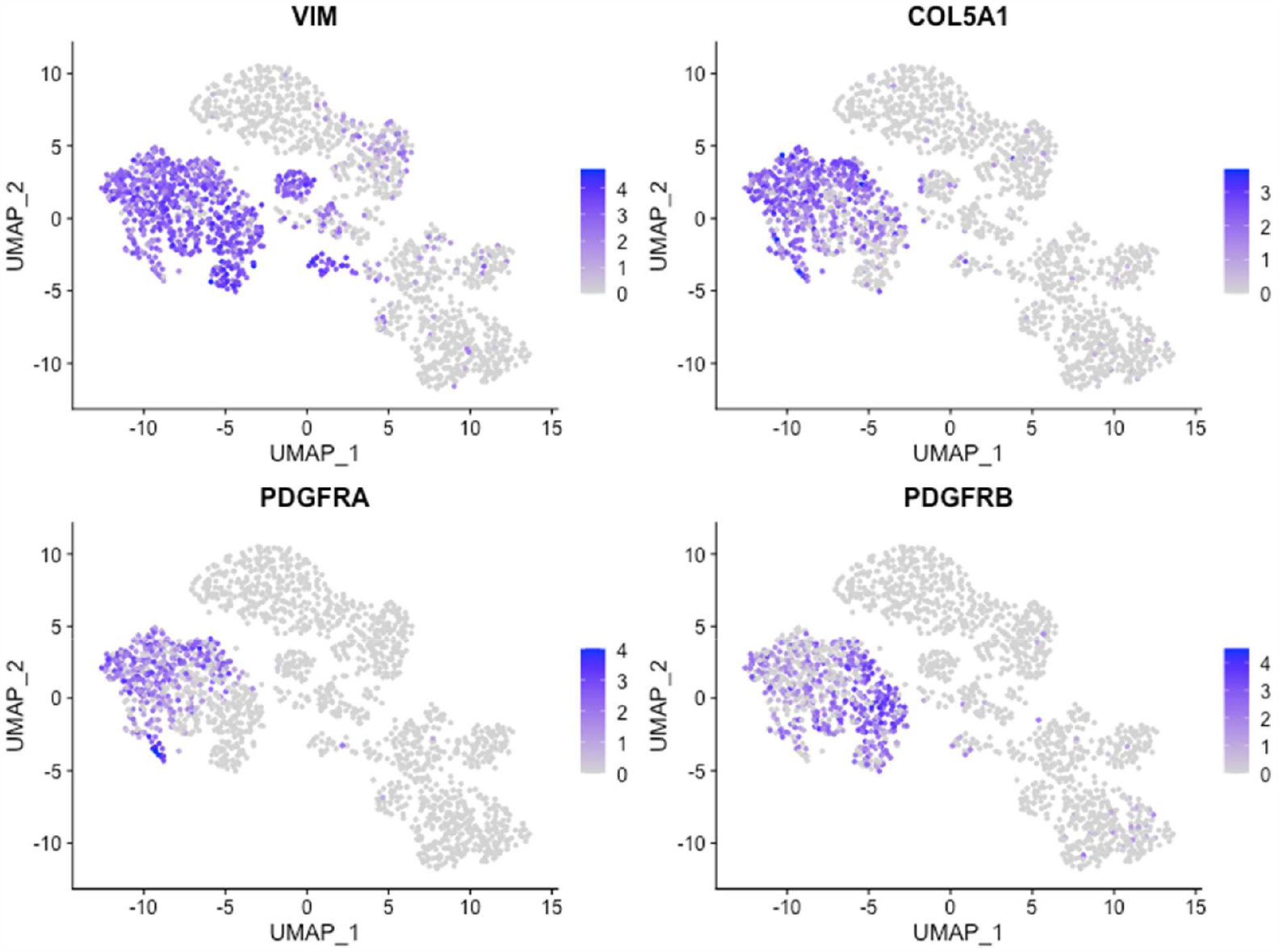
FeaturePlots of gene expression for fibroblast markers (VIM, COL5A1, PDGFRA, PDGFRB)

**Figure 1.2.2.5:**
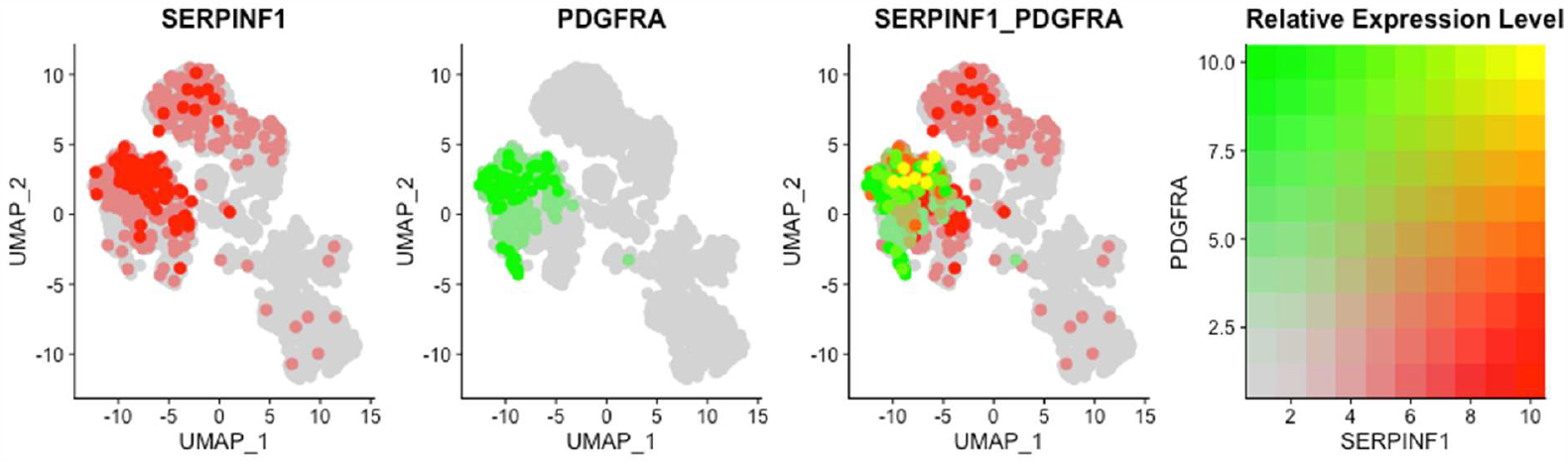
SERPINF1/PEDF and PDGFRA (fibroblast cell marker) gene co-expression FeaturePlots. Far left to far right: Red SERPINF1/PEDF relative gene expression level, Green PDGFRA relative gene expression level, Red SERPINF1/PEDF and green PDGFRA merged (yellow) relative gene expression level, Key of relative gene expression level for preceding three FeaturePlots.

**Figure 1.2.2.6:**
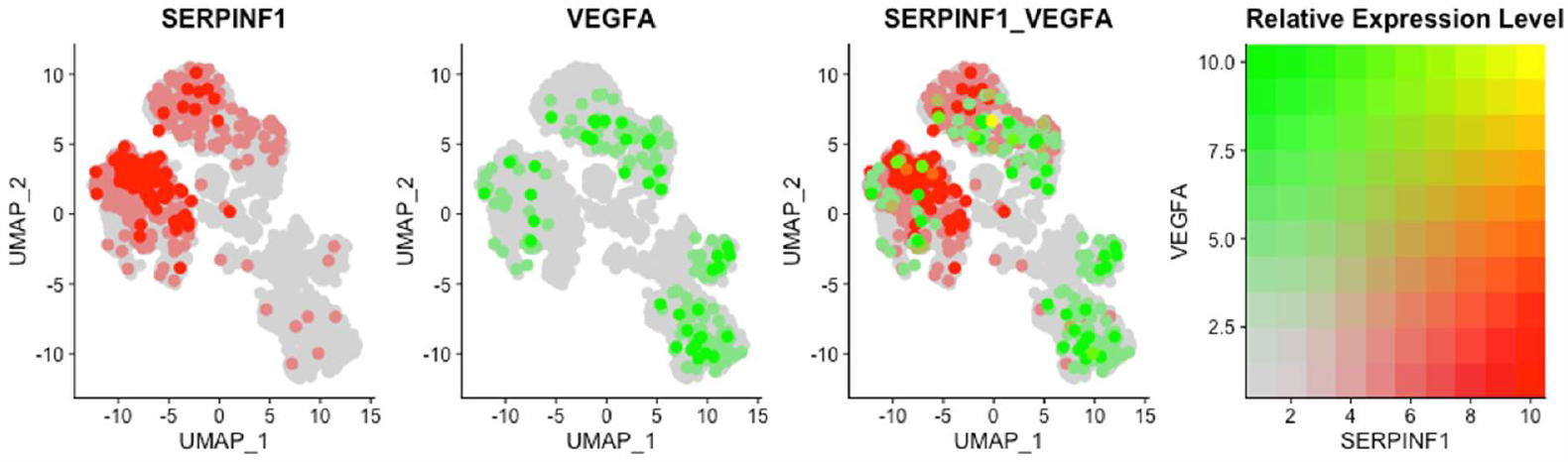
SERPINF1/PEDF and VEGFA (fibroblast cell marker) gene co-expression FeaturePlots. Far left to far right: Red SERPINF1/PEDF relative gene expression level, Green VEGFA relative gene expression level, Red SERPINF1/PEDF and green VEGFA merged (yellow) relative gene expression level, Key of relative gene expression level for preceding three FeaturePlots.

**Figure 1.3.1:**
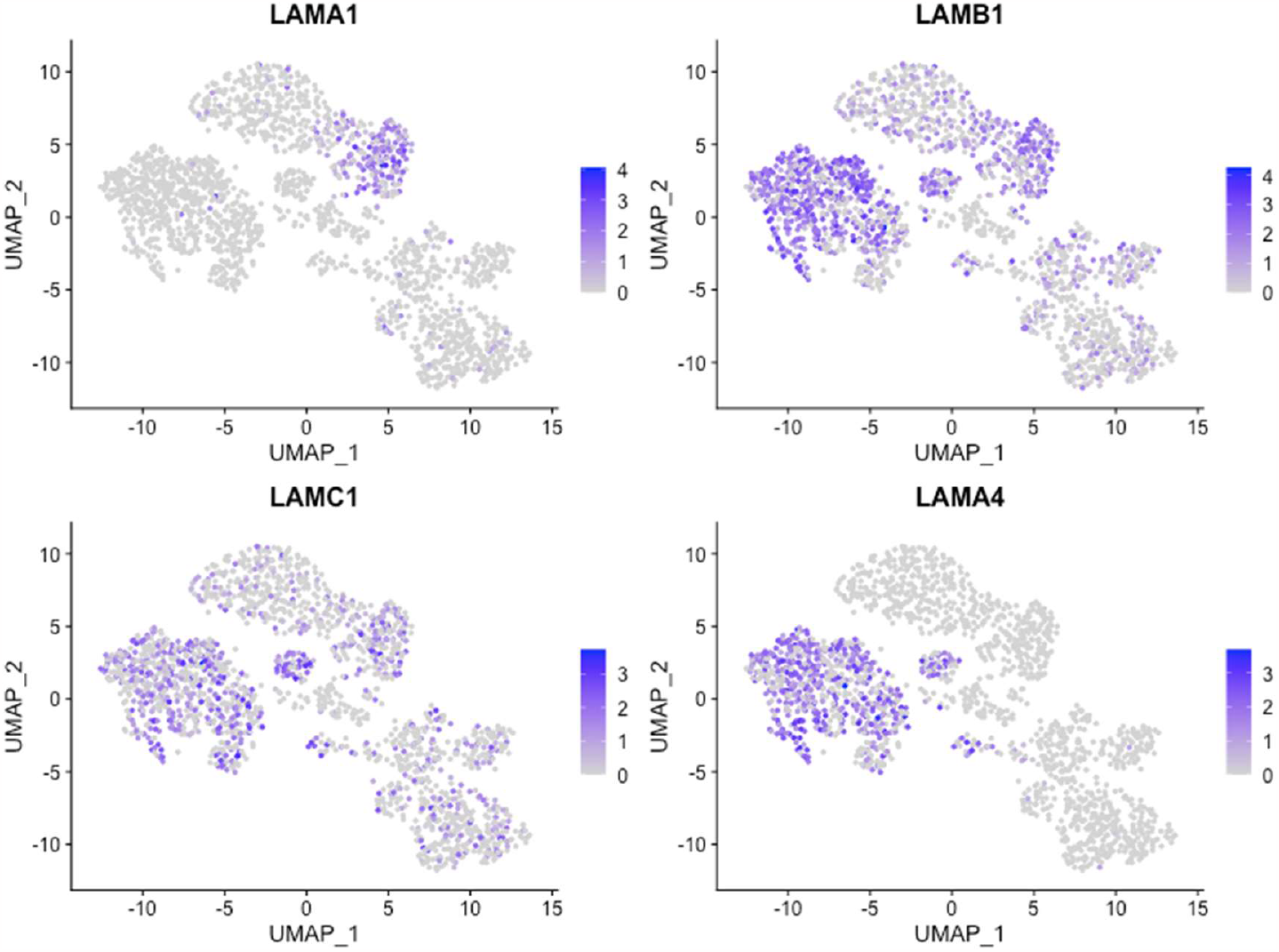
FeaturePlots of gene expression for laminin-α1, laminin-β1, laminin-γ1, and laminin-α4 (LAMA1, LAMB1, LAMC1, LAMA4, respectively)

**Figure 1.3.2:**
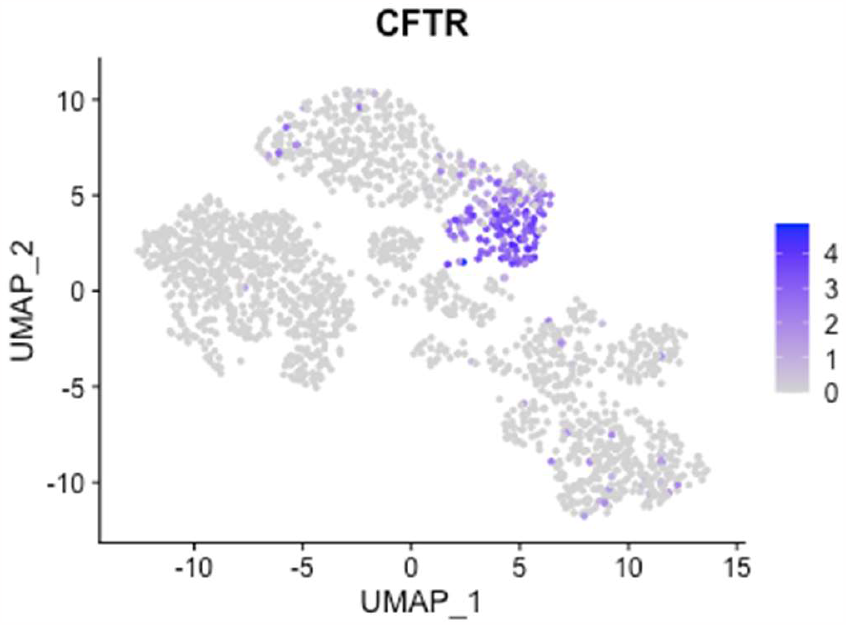
FeaturePlots of gene expression for ductal cell marker (CFTR)

**Figure 1.3.3:**
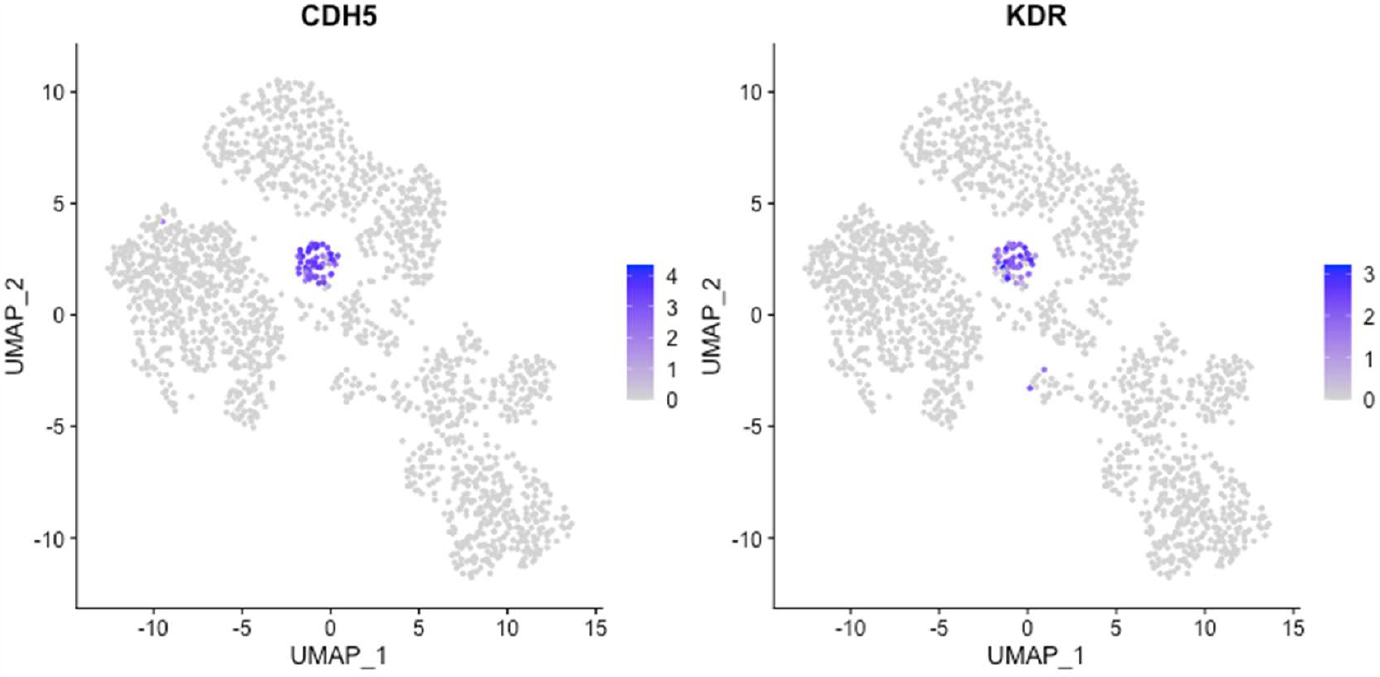
FeaturePlots of gene expression for endothelial cell markers (CDH5 and KDR)

## 1.3 Conclusion

The key finding presented here was the original novel model of pancreatic acinar cells maintaining a microenvironment free from robust angiogenesis by secretion of anti-angiogenic PEDF (SERPINF1). This model has the potential to the bridge a gap in the understanding of human pancreas development.

The existing model that helped initiate the conception of this model is based on the work of Heymans et al^10^. Heymans et al. proposed that endothelial cells promote the deposition, directly or indirectly, of laminin-α1β1γ1 locally around pancreatic endocrine—ductal bipotent (trunk) cells to, essentially, prevent these trunk cells from differentiating into acinar cells.^10^ The trunk is where blood vessels are predominantly localized^10^. Conversely, based on supporting evidence Heymans et al. proposed that the momentary, transient decreased expression or deposition of laminin-α1β1γ1 around pro-acinar tip progenitors allow PTF1-L complex formation^10^. PTF1-L complex formation indicates the transition from pro-acinar tip progenitor cells to differentiated acinar cells.^11^

The simplest combined model suggests SERPINF1/PEDF-secreting pancreatic +PTF1A/+SOX9/+CPA1 pro-acinar tip progenitors and/or SERPINF1/PEDF-secreting pancreatic +COL1A2/+PDGFRA stromal fibroblasts facilitate the momentary, transient decreased expression or deposition of laminin-α1β1γ1 by inhibiting endothelial cells, a major component of blood vessels, by the secreted anti-angiogenic PEDF/SERPINF1 around pro-acinar tip progenitors thereby allowing PTF1-L complex formation.

Interestingly, LAMA1 (laminin α1) was found to be expressed exclusively in ductal cell where as both LAMB1 (laminin β1) and LAMC1 (laminin γ1) was expressed in almost every cell-type as well as endothelial cells with higher gene expression being in the broad mesenchymal cell cluster (Figures 1.3.1, 1.3.2, and 1.3.3). LAMC1 (laminin γ1) overall gene expression was less than LAMB1 (laminin β1) (Figures 1.3.1).

As mentioned in the Introduction (Section 1.1), VEGFA expression in ductal (Figure 1.3.2), endocrine (Figure 1.2.1.1), and mesenchymal cells (Figure 1.2.2.4) supports the model well, because angiogenic VEGFA expression in specifically endocrine cells is expected. Ductal cell VEGFA expression is also somewhat expected because the new model for islet of Langerhans (endocrine cell) generation in the developing pancreas suggests that endocrine cells bud in a peninsula-like manner from the pancreatic bipotent ductal—endocrine trunk.^12^ With endocrine cell generation being intimately connected with the bipotent ductal—endocrine trunk, VEGFA expression in ductal cells is predicted. The model was presented with an additional layer of complexity being that angiogenic VEGFA expression is present in a subset of +PTF1A/+SOX9/+SERPINF1 pro-acinar tip progenitors (Table 1.1). Currently the only reasonable explanation could be that developing acini are not absolutely and completely devoid from vascularization. After all, acini are made up of cells and cells need to exchange nutrition and waste with the blood supply, therefore, there has to be some level of pro-angiogenic regulation occurring within acinar cells. Interestingly, a subset of individual acinar cells were found to be expressing transcripts at relatively high levels for both anti-angiogenic and angiogenic factors simultaneously. (Figure 1.2.2.6)

Overall, this analysis points to the critical impact of angiogenesis regulation in the development of a functional pancreas. This information may be critical for understanding the basic biology of pancreas development and could be useful for developing protocols that generate artificial pancreatic tissue for medical applications and one day the whole organ.

## Notes

https://www.ncbi.nlm.nih.gov/geo/query/acc.cgi?acc=GSE110154

### Competing Interest Statement

The authors have declared no competing interest.

